# The voltage-gated potassium channel *Shal* (K_v_4) contributes to active hearing in *Drosophila*

**DOI:** 10.1101/2024.01.24.577143

**Authors:** Eli S. Gregory, YiFeng Y. J. Xu, Tai-Ting Lee, Mei-ling A. Joiner, Azusa Kamikouchi, Matthew P. Su, Daniel F. Eberl

## Abstract

The full complement of ion channels which influence insect auditory mechanotransduction, and the mechanisms by which their influence is exerted, remain unclear. *Shal* (K_v_4), a *Shaker* family member encoding voltage-gated potassium channels in *Drosophila melanogaster*, has been shown to localize to dendrites in some neuron types, suggesting a potential role for *Shal* in *Drosophila* hearing, including mechanotransduction. A GFP-protein trap was used to visualize the localization of the *Shal* channel in Johnston’s organ neurons responsible for hearing in the antenna. *Shal* protein was localized to the cell body and the proximal dendrite region of sensory neurons, suggesting its involvement not only in general auditory function, but specifically in mechanotransduction. Electrophysiological recordings conducted to assess neural responses to auditory stimuli in mutant *Shal* flies revealed significant decreases in auditory responses. Laser Doppler Vibrometer recordings indicated abnormal antennal free fluctuation frequencies in mutant lines, indicating an effect on active antennal tuning, and thus active transduction mechanisms. This suggests that *Shal* participates in coordinating energy-dependent antennal movements in *Drosophila* that are essential for tuning the antenna to courtship song frequencies.

**Significance Statement:** The study of fruit fly hearing has revealed mechanosensitive ion channels that participate in mechanotransduction, and as in mammalian hearing, energy-dependent mechanisms actively amplify and tune auditory processes. Identifying distinct roles played by different ion channels is essential to better understand this process. Here, we explore the influence of a specific voltage-gated potassium channel, *Shal*, on fly hearing, and find that it affects specific parts of the mechanotransduction process. Our research uncovers *Shal’s* localization in sensory dendrite regions of auditory neurons, where it contributes to shaping mechanotransduction and active antennal tuning. Understanding *Shal*’s involvement in auditory function and mechanotransduction deepens our knowledge of fly hearing and unveils a key player in the coordination of energy-dependent active antennal movements.

## Introduction

Several ion channels participating in insect auditory mechanosensory transduction have been identified, but the precise transduction mechanisms are still poorly understood. For example, the TRPN channel, encoded by the *no mechanotransduction potential C* (*NompC*) (1–3) gene in *Drosophila*, as well as the TRPV channel, comprising two subunits encoded by *inactive* (*iav*) and *nanchung* (*nan*) genes (4–6), are central to mechanotransduction but their precise contributions are yet to be fully established. These channels localize to the sensory cilia of chordotonal organs, also called scolopidia, which make up Johnston’s organ (JO) in the antenna, with NompC localizing distal to the ciliary dilation (7), and Iav/Nan localizing in the ciliary segment proximal to the ciliary dilation (4, 5). Different lines of evidence support two prevailing models of auditory mechanotransduction, known as the NompC model and the Iav/Nan model (8, 9). In the NompC model, TRPN and TRPV act in series, with the NompC channel functioning as the primary mechanotransduction channel to provide the initial transduction current, and Iav/Nan required for propagation as well as providing additional mechanosensitivity. In contrast, the Iav/Nan model attributes mechanosensitivity to Iav/Nan, with NompC acting in parallel to control amplification gain (8–10).

The sensory dendrite, in addition to the being the site of the initial transduction event to generate receptor potentials in response to mechanosensory stimulation, is also involved in active mechanisms. By inserting energy into spontaneous movements of the antenna (2, 11), sensory responses to low amplitude stimulation can be amplified, and antennal movements tuned to enhance reception of sound frequencies related to the *Drosophila* courtship song (12). These active movements are thought to arise from two sources; first, energy representing reduced mechanical compliance upon transduction channel closing, returning kinetic energy into movement of the antenna (13–15); second, active ciliary movement generated by the ATP-dependent axonemal dynein motors in the proximal segment of the sensory cilium (16, 17). It is unclear if and how additional ion channels beyond NompC/Nan/Iav may contribute to the excitability of the sensory cilia in JO, though a number of promising targets exist. This includes the voltage-gated potassium channel encoded by the *Shal* gene, orthologous to K_v_4 (18–20), previously shown in other neurons to localize to dendrites rather than axons (21) and expressed in JO neurons of adult flies (Fly Cell Atlas, 22). If Shal indeed localizes in the dendrites, it has the potential to contribute to shaping JO neuron receptor potential.

Here we first demonstrate that *Shal* is expressed in JO neurons and localizes to the sensory dendrites, including the sensory cilium. Moreover, we show that *Shal* loss of function genotypes result in severe reductions in auditory function, as determined by electrophysiology from the antennal nerve. Furthermore, we found using laser Doppler vibrometry (LDV) that these genotypes disrupt the active mechanisms in JO neurons, especially impairing the active tuning to courtship song frequencies. The K_v_4 channel encoded by *Shal* thus plays a key role in active transduction mechanisms in insect hearing.

## Results

### Shal is expressed in JO neurons and localizes to somata and dendrites

In *Drosophila* single-cell RNAseq data, *Shal* is expressed broadly in JO neurons at the adult stage (22, Fig. S1). A tagged form of *Shal*, when expressed in olfactory projection neurons, has been shown to localize to the soma and dendrites, but not to axons (21). Furthermore, K_v_4.2 in the rat hippocampus has also been found to localize to dendrites (23). Thus, we tested whether Shal channels in JO neurons are localized in the sensory dendrites, where they could contribute to active mechanosensation.

We first used a Shal protein trap line, *Shal^MI00446-GFSTF.1^*, in which GFP and other tags are fused in-frame with the Shal coding sequence in the endogenous locus (24). We found staining for this tagged Shal protein in JO neurons, localized both in the soma and in the dendrites, including in the cilium (Fig. 1A), as indicated relative to the anti-HRP marker for neurons (including dendrites).

**Figure 1.**
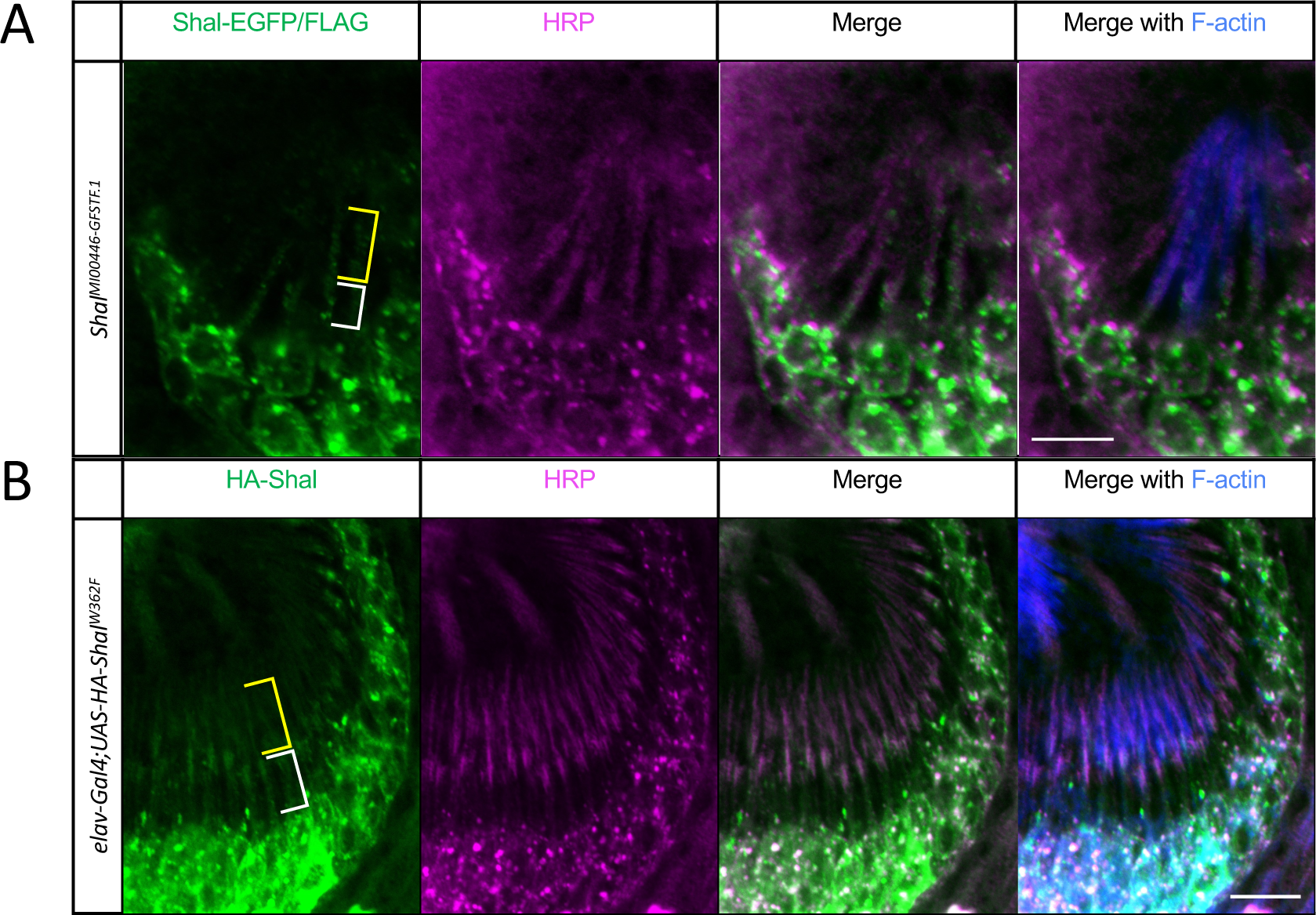
Shal is expressed in JO neurons and localizes to sensory dendrites and somata. A. Immunostaining of pupal JO from the Shal protein trap line, *Shal^MI00446-GFSTF.1^*. The tagged Shal protein is stained with anW-EGFP and anW-FLAG in the green channel. Neurons are visualized with anW-HRP (magenta) which enhances dendrite staining. Phalloidin (blue) stains the scolopale rods in the scolopale cell surrounding the sensory dendrite. Brackets indicate the inner (blue bracket) and outer (orange bracke) dendriWc segments. Scale bar = 5 μm. B. Immunostaining of pupal JO from flies expressing the Shal dominant-negaWve construct, *UAS*519 *HA-Shal^W362F^*, in neurons. AnW-HA staining shows the dominant-negaWve construct in green, with anW-HRP (magenta) and phalloidin (blue). Brackets as in A. Scale bar = 10 μm.

To confirm this localization, we also used an HA-tagged dominant-negative construct (Table 1, 19) expressed in all neurons with the *elav^C155^-Gal4* driver. Anti-HA staining in this genotype also showed staining in the dendrites co-linear with the anti-HRP marker (Fig. 1B). Thus, the expression pattern of Shal suggests a physiological role in JO neurons, and its localization to dendrites is consistent with a possible role in sensory transduction mechanisms.

**Table 1.**
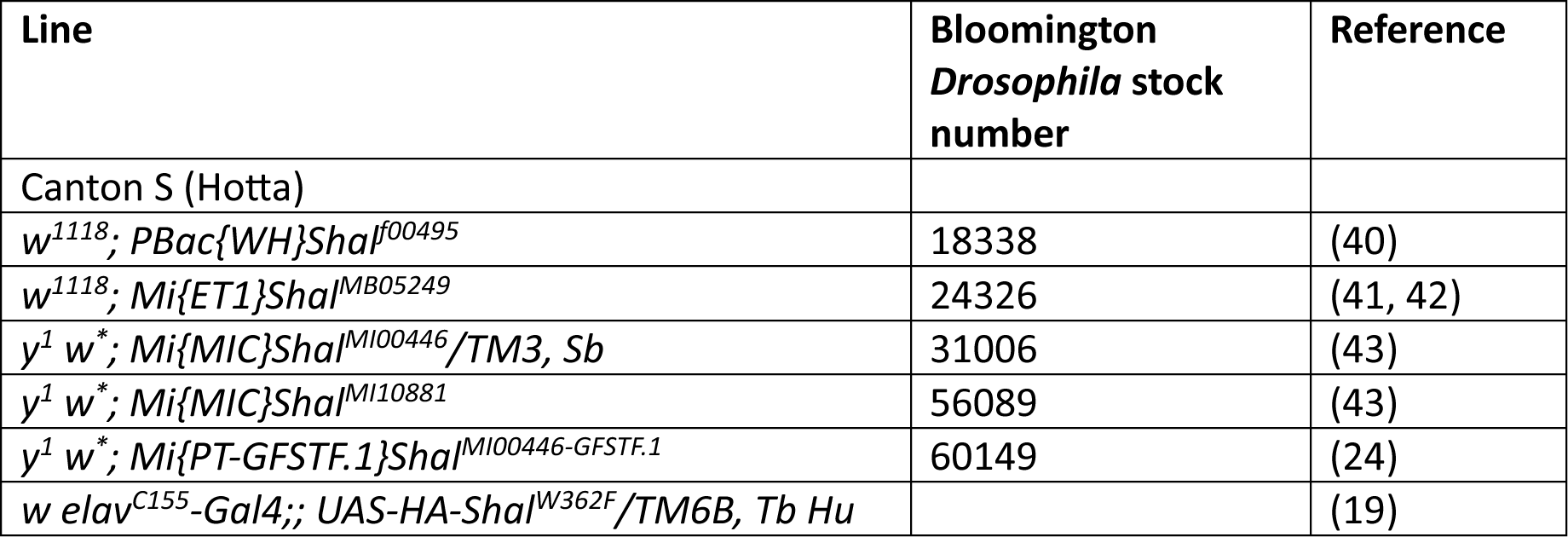
*Drosophila* genotypes used in this study.

### Shal is required for proper auditory function in Drosophila

To test whether auditory function was impaired following *Shal* mutation, we tested several *Shal* loss of function genotypes (Table 1, Fig. S2) for changes in sound-evoked potentials (SEPs) (25, 26). In this electrophysiological assay, field potentials recorded from the antennal nerve at the joint between segments 1 and 2 represent the combined auditory signals in the axons from all JO neurons (Fig. 2A).

**Figure 2.**
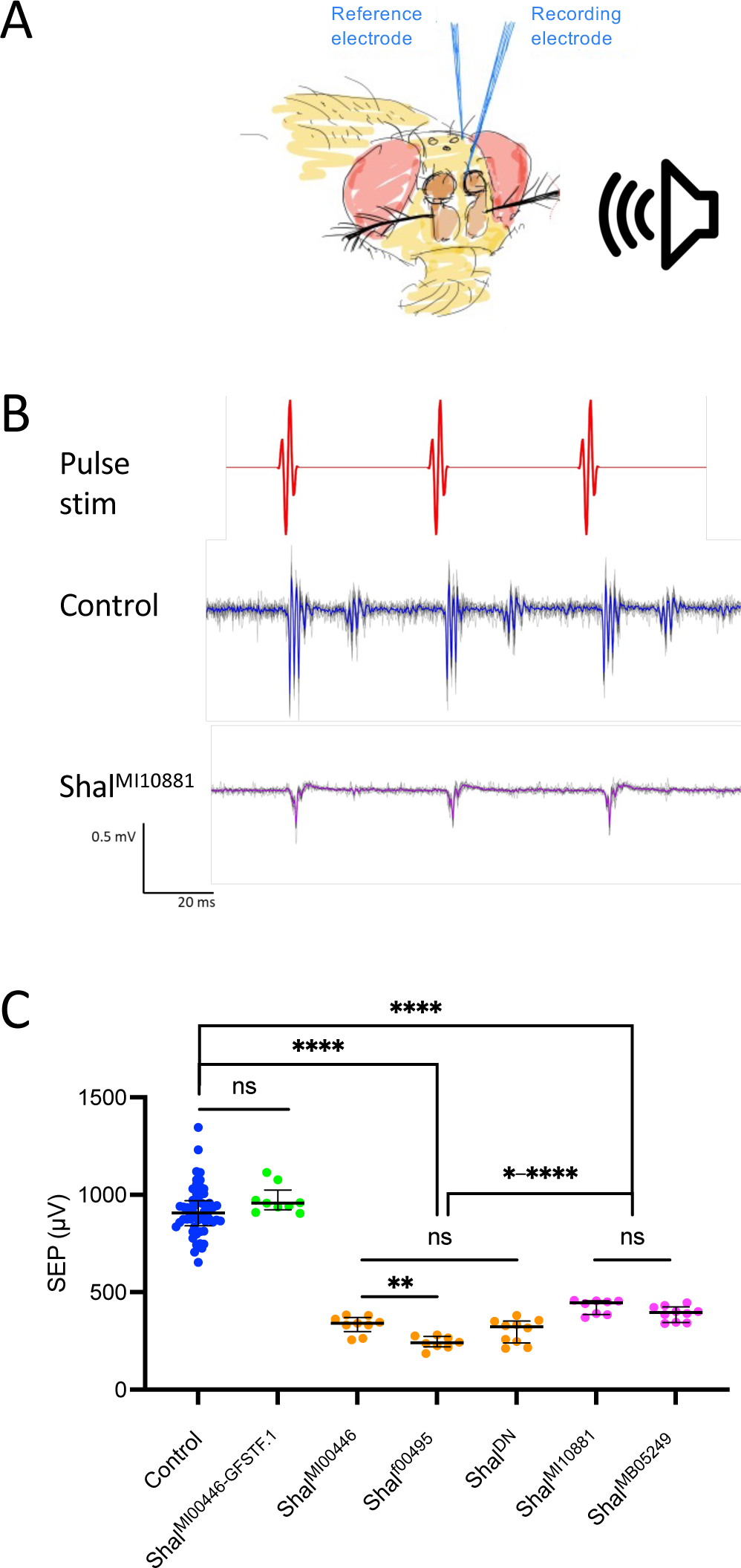
Shal loss of function impairs auditory signals in the antennal nerve. A. Schematic of electrophysiological recording preparation. Sound-evoked potentials (SEPs) are recorded as differential field potentials from a tungsten electrode placed near the antennal nerve (recording electrode) relative to one inserted in the dorsal head capsule (reference electrode) in response to presentation of near-field acoustic stimuli. B. Example traces from control and mutant flies in response to synthetic pulse song stimulus. Individual responses to ten consecutive stimuli are depicted as thin gray lines, and their average as the thicker blue (control) or magenta (mutant) line, in agreement with the color scheme in C. C. Scatter plot of SEPs recorded from flies with *Shal*-related genotypes. Each dot represents the SEP amplitude recorded from one antenna. Bars indicate means and error bars represent SEM. Controls (blue dots) and the Shal protein trap flies, *Shal^MI00446-GFSTF.1^*(green dots) are not significantly different, but all other genotypes are significantly different from controls. Strong alleles (orange dots) produce significantly lower SEPs than weak alleles (magenta dots). The dominant-negative Shal genotype (ShalDN = *w elav^C155^-Gal4;; UAS-HA-Shal^W362F^/TM6B, Tb Hu*) behaves as a strong mutant genotype. Brown-Forsythe ANOVA, p < 0.0001, with Dunnett’s multiple comparisons (ns: not significant; *p < 0.05; **p < 0.01; ***p < 0.001; ****p < 0.0001).

In contrast to controls, *Shal* insertion mutants all showed significantly reduced SEP amplitudes (Fig. 2B,C; ANOVA(Brown-Forsythe); p < 0.0001). Compared to controls, two insertion alleles, *Shal^MI00446^ and Shal^f00495^*, showed the most reduced SEPs (Fig. 2C; Dunnett’s T3 multiple comparisons, p < 0.0001 each), while *Shal^MI10881^* and *Shal^MB05249^* showed SEPs that were significantly reduced compared to controls (Dunnett’s; p < 0.0001), but significantly higher than the other two alleles (Fig. 2C; Dunnett’s; p values between 0.006 and < 0.0001). To test for possible changes in hearing function, we also tested the protein trap line *Shal^MI00446-GFSTF.1^*, derived from *Shal^MI00446^* by replacing the MiMIC cassette in the insertion with the GFSTF marker cassette flanked by splice acceptor and splice donor sites (Table 1, Fig. S2) to generate full length *Shal* proteins with the markers fused in frame (24). SEPs of this protein trap line were not significantly different from controls (Dunnett’s; p = 0.60), and significantly different from the parent *Shal^MI00446^* allele (Fig. 2C; Dunnett’s; p < 0.0001), suggesting that the fusion protein is fully functional.

Finally, flies expressing the Shal dominant-negative construct in all neurons also showed strong reduction in SEPs compared to controls, (Fig. 2C; Dunnett’s, p < 0.0001). In fact the Shal dominant-negative SEPs were not significantly different from the stronger insertion alleles, *Shal^MI00446^ and Shal^f00495^* (Fig. 2C; Dunnett’s, p = 0.97 and p = 0.28, respectively), but significantly more reduced than the two weaker insertion alleles, *Shal^MI10881^* and *Shal^MB05249^* (Fig. 2C; Dunnett’s; p = 0.0014 and p = 0.0223, respectively). In summary, all mutant or dominant-negative genotypes resulted in significant SEP reductions, and the protein trap fusion restored function to the mutant insertion from which it was derived. All together this indicates that *Shal* plays critical roles in sending auditory signals to the brain.

### Shal is required for tuning antennal active movements

To test whether Shal channel activity in the dendrite is important for active mechanosensation to tune the antenna’s resonant frequency and energy injection into antennal movement, we measured free fluctuations (FF) of *Shal* mutant and control fly antennae, both awake and under CO_2_ sedation, using LDV (2, 11, 27) (Fig. 3A). In awake flies, we found significant differences in resonant frequency based on genotype (Brown-Forsythe ANOVA, p < 0.0001). The antennal resonant frequency of awake control flies was 240.9 ± 9.1 Hz (mean ± SEM; Fig. 3B). Among *Shal* insertion mutants, *Shal^MB05249^* and *Shal^MI10881^* flies showed no significant change in awake tuning, with 260.6 ± 27.8 Hz and 197.0 ± 24.7 Hz respectively (Fig. 3B) (Dunnett’s T3 multiple comparisons, p > 0.9999 and p = 0.84, respectively). In contrast, *Shal^f00495^* and *Shal^MI00446^* flies showed significant increases in the awake tuning, with 423.7 ± 15.8 Hz and 340.9 ± 8.8 Hz respectively (Fig. 3B) (Dunnett’s p < 0.0001 each). Furthermore, the dominant-negative *Shal* flies also showed increased awake tuning of 338.9 ± 23.6 Hz (Fig. 3B) (Dunnett’s p < 0.033). Meanwhile, the *Shal* protein trap flies were not distinguishable from controls, at 273.3 ± 19.4 Hz (Dunnett’s p = 0.091).

**Figure 3.**
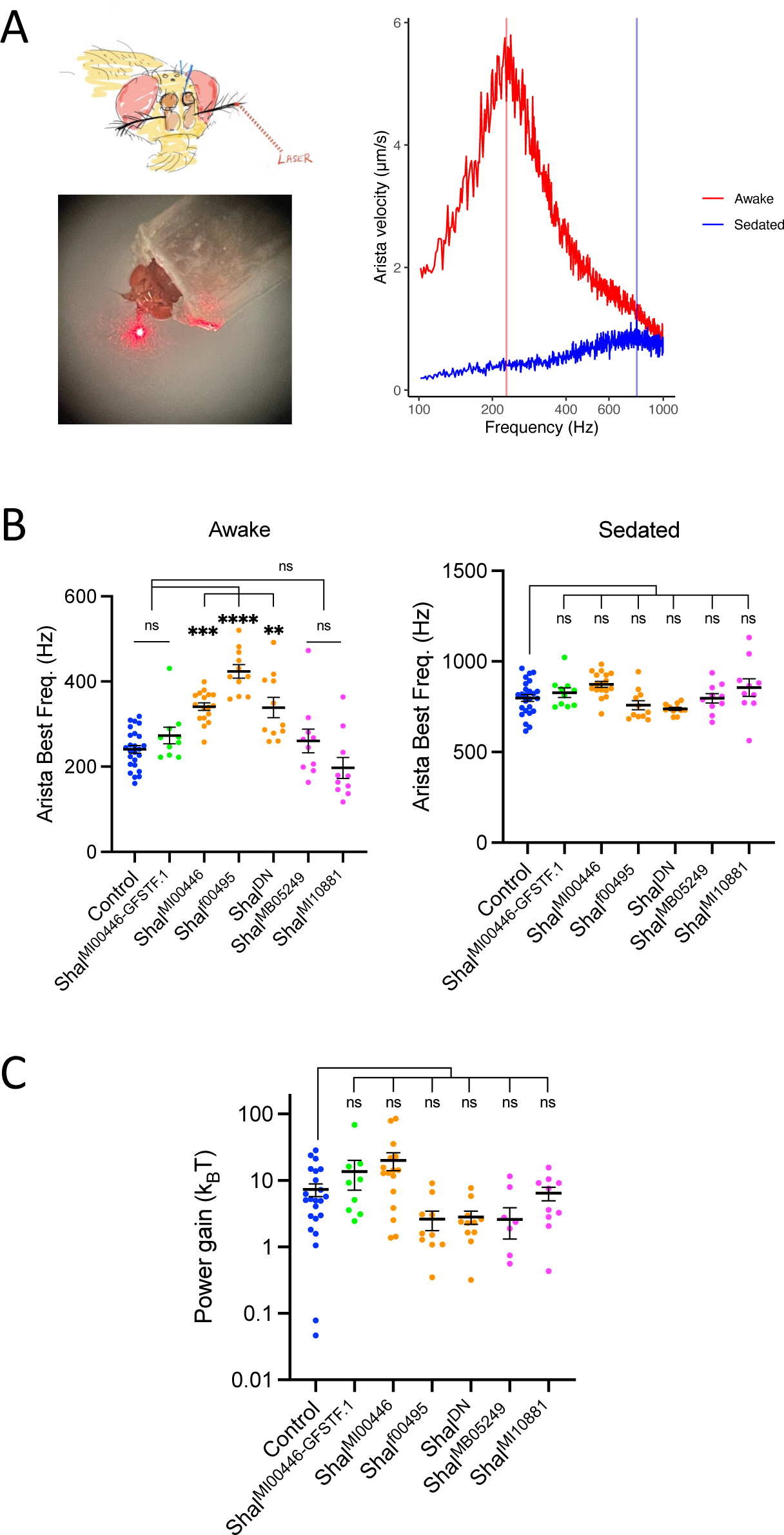
Shal loss of function shifts antennal resonant frequency with little effect on power gain. A. Laser Doppler vibrometry preparation. Reflections from a laser beam focused on the arista allow precise recording of antennal movements. In the awake state (red trace), the antenna of a control fly shows vibrations in a range of frequencies below 1 kHz, with a peak at about 240 Hz (vertical red line). The same antenna under CO_2_ sedation (blue trace) shows lower magnitude vibrations with a peak in the 800 Hz range (blue vertical line). These recordings in the absence of sound stimuli are called “free fluctuations”. B. Scatter plots of the peaks (best frequencies) of antennal free fluctuations in the awake state (left graph) and the sedated state (right graph). Each dot represents the best frequency on one antenna. Bars indicate means and error bars represent SEM. Colors of dots match genotypes of Fig. 2. In the awake state, the strong alleles (orange dots) show best frequencies significantly higher than controls (blue dots). However, weak alleles (magenta dots) do not significantly shift the best frequencies compared to controls. In the sedated state, none of the genotypes significantly differ from controls. Brown-Forsythe ANOVA, p < 0.0001, with Dunnett’s multiple comparisons (ns: not significant; **p < 0.01; ***p < 0.001; ****p < 0.0001). C. Scatter plot of estimated power gain calculations. Genotypes and dot colors as in B. Power gains in Shal mutant genotypes do not differ significantly from controls. Kruskal-Wallis ANOVA with Dunn’s multiple comparisons (ns: not significant).

**Figure 4.**
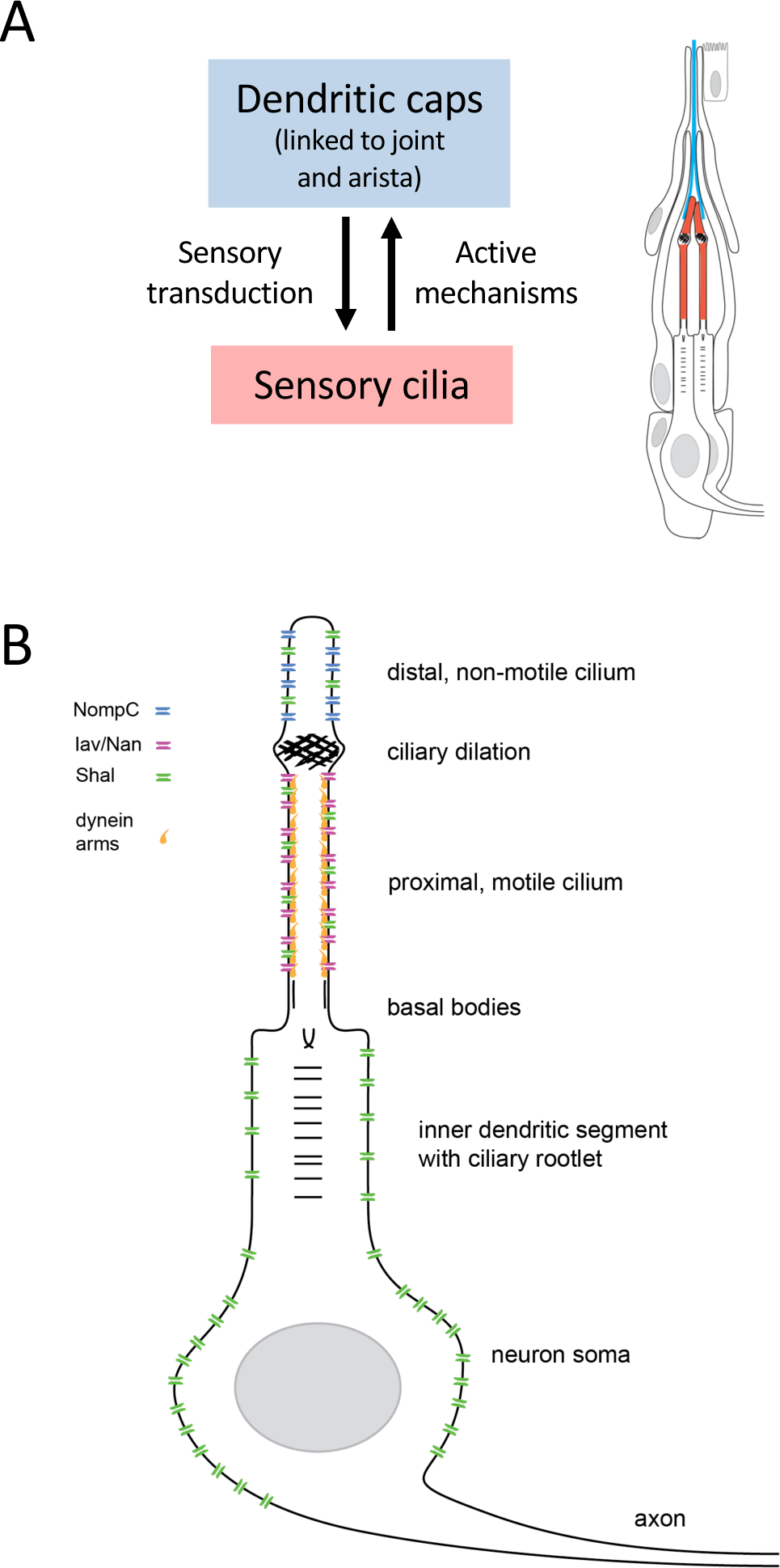
Model of JO neuron ion channels in active mechanotransduction. A. During sensory transduction, sound-induced movements of the arista are transferred by the dendritic caps (blue) to the JO neuron ciliated dendrites (red) to initiate mechanotransduction. In the absence of sound stimuli, active ciliary movements in the sensory dendrites of JO neurons transfer kinetic energy to the antennal joint via the dendritic cap result in antennal vibrations (free fluctuations). B. In the JO neurons, the several ion channels have been localized to the dendritic compartment. NompC (TRPN) is localized in the distal-most ciliary compartment beyond the ciliary dilation. This ciliary segment is non-motile given the absence of axonemal dynein arms. Nan/Iav (TRPV) channels localize in the motile proximal ciliary segment, co-linear with the localization of axonemal dynein arms. In this study, we show that Shal localizes along the entire sensory cilium where it can likely modulate the dynamics of currents mediated by both TRPN and TRPV and thereby impact the timing of active ciliary movements. Shal also localizes in the neuron soma and in the inner dendritic segment connecting the cilium and the soma. In these regions, Shal may affect the propagation of the sensory receptor potential or its conversion into an action potential. Some alleles may affect this somatic function, disrupting the generation or propagation of full action potentials without affecting active ciliary movements.

Sedation with CO_2_ removes physiologically active process, leaving only passive mechanical movements of the antenna (Fig. 3A) (2, 11, 27). Under sedation, we found the resonant frequency of control flies to be 797.9 ± 18.8 Hz (Fig. 3C), as expected from previous reports (12). We also found all *Shal* genotypes tested to show passive resonant frequencies in the same range (Fig. 3C) and not significantly different from controls (Dunnett’s multiple comparisons p-values ranging from p = 0.087 to p > 0.9999), although the overall Brown-Forsythe ANOVA was significant at p = 0.0058, primarily attributable to a difference between *Shal^MI00446^* and the Shal dominant-negative flies. These findings show that Shal has minimal impact on passive mechanical properties of the antenna, but is required to shift the antennal tuning from the passive 800 Hz range to the fully active 240 Hz range. Without Shal activity in the strongest loss of function genotypes, the active tuning shifts only partially, to the 400 Hz range.

From the active and passive LDV recordings, it is possible to calculate auditory power gain, the active injection of energy into the hearing system representing one measure of the energy provided by the active system above the passive baseline. While a Kruskall-Wallis ANOVA test of power gain calculations (Fig. 3C) shows significant differences by genotype (p = 0.0015), none of the individual genotypes is significantly different from controls (Dunn’s multiple comparisons p values range from p = 0.728 to p > 0.9999). Significance in the overall model arise primarily from *Shal^MI00446^* vs. *Shal^f00495^* (Dunn’s p = 0.02) and *Shal^MI00446^* vs. *Shal^MB05249^* (Dunn’s p = 0.011). Compared to controls, we see no obvious changes in power gain among *Shal* loss of function genotypes, suggesting that Shal has little effect on the overall energy that JO neurons generate for active mechanosensation.

## Discussion

We have shown that Shal is expressed in JO neurons where it is important for the output of these neurons, as measured by the electrophysiological signals sent to the brain along the antennal nerve. Shal localizes not only to the JO neuron cell bodies but also to the sensory dendrite where it is positioned to participate in active hearing mechanisms. Indeed, our LDV data confirm that loss of Shal significantly affects the active physiological tuning of antennal oscillation in the absence of sound. In wild-type flies, such tuning mechanisms shift antennal oscillations from a resonant frequency in the 800 Hz range to the 240 Hz range.

We were surprised to find that some alleles, specifically *Shal^MI10881^* and *Shal^MB05249^*, reduced SEPs but had little effect on the active mechanisms of antennal movement. This suggests that Shal may have distinct contributions to these active mechanisms, presumably acting in the dendrite, compared to the generation or propagation of the action potential measured in the nerve, functions that may depend more on Shal channels localized in the soma. Thus, these two alleles may differ in the specific properties of the Shal channel required for localization to these two compartments or of the Shal channel K^+^ currents as they contribute to these different functions. Alternatively, these two functions may simply differ in their susceptibility to reduced expression levels. Interestingly, the *Shal^MI10881^* and *Shal^MB05249^* alleles are inserted into the first intron, in the 5’UTR region, while the other insertion alleles are inserted in the second intron, located in the coding region (Fig. S2). This difference in insertion site may allow any read-through transcripts still to generate normal full-length Shal channel proteins for those insertions in the first intron, albeit at reduced levels, while read-through transcripts from those in the second intron will affect the channel structure.

The dominant-negative Shal is associated with a single amino acid change, from tryptophan to phenylamine at position 362 (19), located at the pore region, based on a similar mutation in mouse K_v_4.2 (28). K_v_4 channels assemble as tetramers (29), and incorporation of K_v_4.2^W362F^ subunits has been shown to block channel activity (28). Overexpression of *UAS-HA-Shal^W362F^* in the background of normal *Shal* alleles is likely to result in a preponderance of mutant channel subunits, making assembly of a wild-type tetramer rare and leading to a dominant-negative phenotype equivalent to a strong loss of function phenotype. This interpretation is consistent with our results in both the electrophysiological and antennal movement phenotypes (Fig. 2C, Fig. 3A).

The cellular trafficking mechanisms that localize the Shal protein in JO neuron dendrites are unknown. In olfactory projection neurons, dendritic localization of Shal depends on a di-leucine motif and its interacting protein, Shal-interacting Di-leucine protein (SIDL) (21). The Shal locus has been shown to generate three distinct mRNA splicing isoforms which differ in the 3’ end (18; Fig. S2, 30). The di-leucine is located at residues 481-482, about 7 residues before the divergence of the three isoforms. Thus, all three isoforms contain the di-leucine motif, but as the dendritic targeting was tested in the context of the longest isoform, and it is not known which isoforms are expressed in JO neurons, we cannot be certain whether this dendritic targeting mechanism applies to Shal in JO neurons. Furthermore, while *SIDL* is expressed in JO neurons, its expression level is low (Fly Cell Atlas, 22). K_v_4.2 and K_v_4.3 localization in the rat hippocampus may rely on multiple mechanisms, including localization of the mRNA based on sequences in the 3’-UTR or protein localization through K_v_ channel interacting proteins (KChIPs) or membrane-spanning dipeptidyl aminopeptidase-like proteins (DPPs) (23, 31–33). Whether any of these mechanisms play a role in JO neurons remains to be tested.

The specific mechanism by which dendritic Shal channels shape antennal motion is not known. Active antennal motion is presumably performed by axonemal dyneins (16, 17), and their regulation must require appropriate membrane potential dynamics to coordinate the ciliary movement in the correct phase to change the frequency appropriately. Our findings of minimal effect on power gain in *Shal* mutants (Fig 3C), with effects primarily on tuning, suggest the strength of ciliary activity is largely unaffected, but is rather out of phase with membrane potential changes, and inhibiting tuning to the desired 240 Hz range.

One possible mechanism by which Shal channels might affect receptor potential dynamics locally in the dendrite is to sharpen receptor potentials through their fast activation and inactivation kinetics (18, 30, 34, 35). In hippocampal CA1 pyramidal cell dendrites and granule cell dendrites in the dentate gyrus, the transient A-type potassium currents (encoded by K_v_4) have also been reported to inhibit backpropagation of action potentials, limit the dendritic initiation of action potentials and dampen the effect of excitatory dendritic inputs (36–38). Thus, in JO neuron dendrites, Shal may be shaping the kinetics of one receptor potential as it develops, and may also modulate any back propagation to coordinate with the next cycle of receptor potential activation to sustain active oscillations. These activities likely also depend on other dendritic ion channels, including NompC and Nan/Iav, as well as potentially the voltage-gated sodium channel encoded by the *para* gene, which has also been shown to localize to JO neuron dendrites (39). Detailed understanding of how all these channel activities interact to shape receptor potentials and modulate the timing of axonemal dynein arm activity to influence antennal motion will require further investigation.

## Materials and Methods

### Fly strains

*Drosophila melanogaster* genotypes used in this study are listed in Table 1. Controls included the wild-type Canton S strain, as well as TM3, Sb heterozygotes from the Shal^MI00446^ strain, and in some experiments, *w elav^C155^;; TM6B, Tb Hu/+* flies resulting from outcrossing the balanced dominant-negative flies listed in the table with a cantonized *w^1118^* strain. In all experiments, the different controls were tested to ensure no significant differences before pooling. For all experiments, *Shal^MI00446^* flies were homozygotes selected from the balanced stock, and dominant-negative Shal flies were of the genotype shown in the table.

### Antibody staining and imaging

*Drosophila* pupal heads were dissected in cold phosphate-buffered saline (PBS), then fixed in 4% paraformaldehyde (PFA) for 15 minutes. After three washes in PBS with 0.2% Tween-20 (PBT) with rotation over 30 minutes, the samples were blocked with Blocking Buffer (BB; freshwater fish skin gelatin, normal goat serum and bovine serum albumin in PBT) for 1 hour with rotation. Primary antibodies diluted in BB included mouse anti-FLAG antibody (1:100; Sigma), mouse anti-GFP antibody (1:500; Thermo Fisher Scientific), anti-HA rMs-IgG1 (1:500; DSHB) and rabbit anti-HRP antibody (1:500; Cappel). Primary antibody incubation took place overnight at 4°C with rotation.

The following day, the samples were washed three times in PBT over 30 minutes with rotation, then incubated with secondary antibodies and phalloidin for 2 hours at room temperature with rotation. Secondary antibody diluted in BB (1:500) consisted of goat anti-mouse Oregon Green (488; Thermo Fisher Scientific) and goat anti-rabbit TRITC (Jackson Immunoresearch), along with phalloidin 405 (1:1000; Thermo Fisher Scientific). Samples underwent two 15-minute washes in PBT with rotation followed by a brief 5-minute wash in PBS. Finally, samples were mounted in Fluoromount (Thermo Fisher Scientific) onto glass slides with 1.5 coverslips and imaged with a Leica STELLARIS 8 confocal microscope, using a 63x objective lens with oil immersion.

### Electrophysiology

Sound-evoked potentials (SEPs) were captured using a pair of electrolytically sharpened tungsten recording electrodes (25, 26). The recording electrode was inserted between the first and secondary segments of the antennae, while the reference electrode was inserted into the head cuticle near the posterior orbital bristle. A computer-generated pulse song was introduced frontally to the fly under near-field conditions.

Signals were subtracted and amplified with a differential amplifier (DAM50, World Precision Instruments) and digitized at 10 kHz (USB-6001, National Instruments). Average response values were measured as the max-min values in an averaged trace from 10 consecutive presentations of the described protocol. SEP data were analyzed using one-way ANOVA multiple comparisons with Welch’s correction for unequal variances.

### Laser Doppler Vibrometry data collection

Following 2 days of entrainment in a 12:12 LD regime, male and female *Drosophila* aged between 3 and 10 days old were aspirated into micropipette tips as for electrophysiology experiments. Fly head movement was restricted by the application of modelling clay to the edge of the pipette. Blue light-cured glue was then applied to the entirety of the right antennae (to completely inhibit movement) as well as the base of the left antennae.

The tip with the immobilized fly was then attached to a rod held in a micromanipulator on a vibration isolation table in a temperature-controlled room (set to 25°C ± 1°C). The fly was positioned such that its left arista was perpendicular to the beam of a laser Doppler vibrometer (Vibroflex, Polytec).

Unstimulated aristal vibrations (denoted as ‘free fluctuations’) were first recorded while the fly was awake. The fly was then sedated via continuous CO_2_ exposure for 2 minutes, before another free fluctuation recording was made. These recordings allowed for investigating both active (awake) and passive (sedated) hearing states.

### Laser Doppler Vibrometry data analysis: mechanical tuning calculation

Fast Fourier transforms of recording values were made using the Vibsoft Polytec software for frequencies from 1 Hz to 10 kHz. Frequency values below 100 Hz were excluded from analyses due to significant noise in the recordings.

A forced damped oscillator function was applied to transformed data via the lme4 package (version 1.1-33) in R (version 4.3.0):

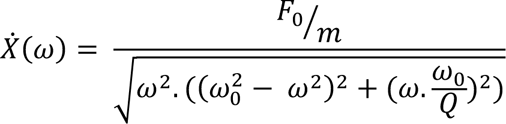

Here, F_0_ = external force strength

m = flagellar apparent mass

*ω* = angular frequency

*ω*_0_ = natural angular frequency

Q = quality factor = m*ω*0/*γ* (*γ* = damping constant).

Application of this function allowed for estimation of the natural angular frequency of recordings from both active and passive states for each fly, enabling calculation of the mechanical tuning frequency, f_0_ (= ω_0_/2π).

### Laser Doppler Vibrometry data analysis: power gain calculation

Power gain calculations utilized the results of the forced damped oscillator function fits by enabling the calculation of the ratio of total fluctuation power of an individual’s active and passive states.

We defined power gain as:

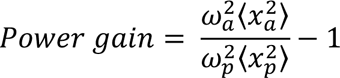

Here,

*ω*_*a*_= natural angular frequency of active system

*ω*_*p*_= natural angular frequency of passive system

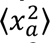 = sum of squared Fourier displacement amplitudes in active state

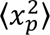= sum of squared Fourier displacement amplitudes in passive state

Natural angular frequency values were calculated from the function fits, while sums of squared Fourier displacement amplitudes were estimated from:

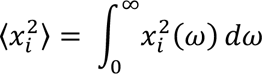

## Acknowledgements

We are grateful to Susan Tsunoda for providing flies with the Shal dominant negative construct. We also thank the Bloomington *Drosophila* Stock Center for fly stocks. Monoclonal antibodies were obtained from the Developmental Studies Hybridoma Bank, created by the NICHD of the NIH and maintained at The University of Iowa, Department of Biology, Iowa City, IA 52242. Funding is acknowledged from NSF grant 2037828 (to Alan Kay, DFE, and Zahre Aminzare), Iowa Office for Undergraduate Research (ESG), JSPS Invitational Fellowships for Research in Japan (Short-term) (S22091 to DFE), The Nagoya University Neuroscience Institute (DFE), International Principal Investigator (PI) Invitation Program, Nagoya University, Japan (DFE), Tokai Pathways to Global Excellence, Nagoya University, Japan (0121an0002 to MPS).

## CRediT Roles

Conceptualization MPS, DFE

Data Curation ESG, YYJX, TTL, MAJ, MPS, DFE

Formal Analysis ESG, YYJX, TTL, MAJ, MPS, DFE

Funding Acquisition AK, MPS, DFE

Investigation ESG, YYJX, TTL, MAJ, MPS, DFE

Methodology ESG, MAJ, MPS, DFE

Project Administration AK, MPS, DFE

Resources AK, MPS, DFE

Software MPS, DFE

Supervision AK, MPS, DFE

Validation ESG, YYJX, MAJ, MPS, DFE

Visualization ESG, MAJ, DFE

Writing – Original Draft ESG, MPS, DFE

Writing – Review and Editing ESG, YYJX, AK, MPS, DFE

