## Supplemental Figures S1 and S2 for "The voltage-gated potassium channel *Shal* (K_v_4) contributes to active hearing in *Drosophila*"

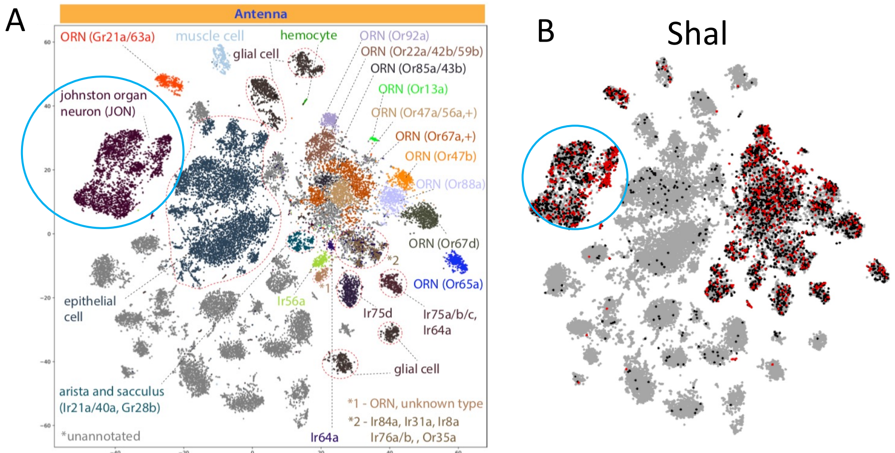


**Fig. S1. Expression of Shal in antennal single nucleus RNA sequencing (Fly Cell Atlas)**

A. Annotated clustering of single-nucleus RNA transcript expression from antenna (reproduced from Li et al (2022) with permission), showing a cluster of cells representing the JO neurons (circled). B. Expression of Shal (red) depicted over the same clusters indicates that Shal is expressed in JO neurons (circled) as well as olfactory neurons.

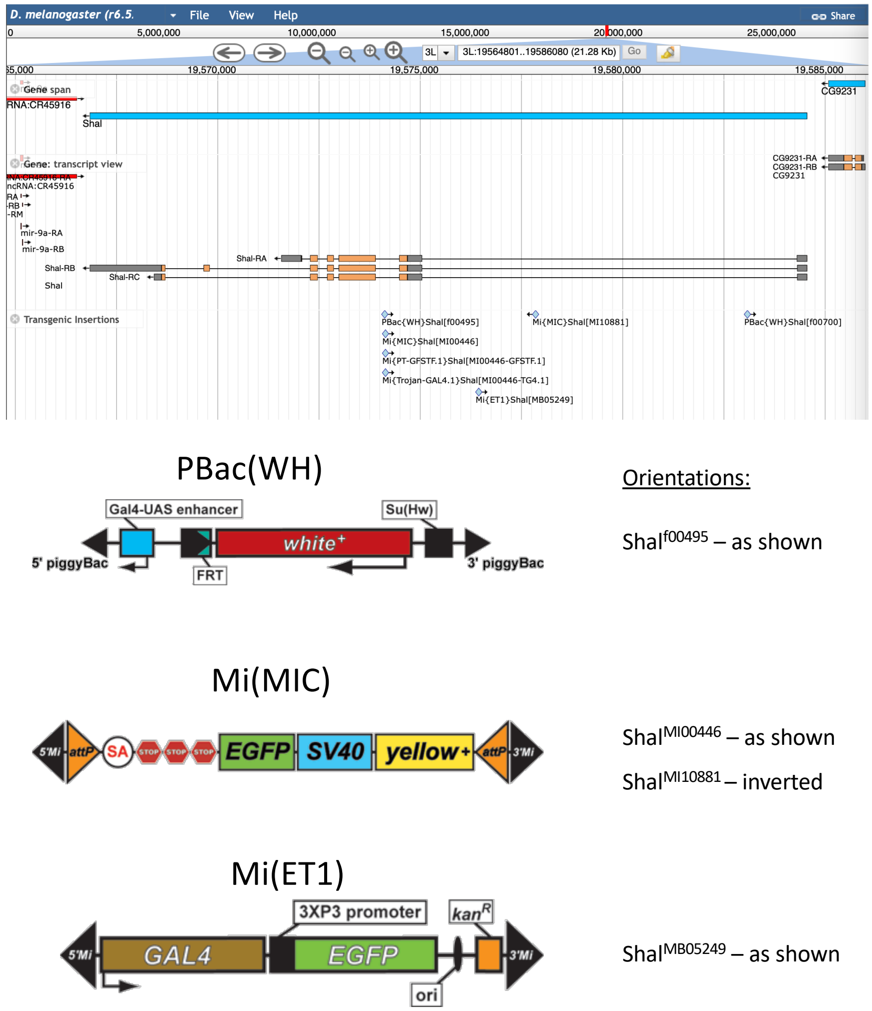


**Fig. S2. Map of *Shal* locus.**

Upper panel shows a screenshot of the JBrowse genome browser depicting the *Shal* locus on chromosome 3L. *Shal* is transcribed in the leftward direction, with three transcript splice isoforms (coding regions in orange boxes, non-coding regions in gray). Transposon insertion sites are depicted by small blue triangles, labeled. Corresponding transposon structures are diagrammed below (from the Gene Disruption Project (https://flypush.research.bcm.edu/pscreen/transposons.html)), with orientation information relative to the map.
